# Cryo-EM structural analysis of human TOP1 trapping by eight clinical anticancer drugs

**DOI:** 10.64898/2026.08.03.742490

**Authors:** Xi Yang, Yves Pommier

## Abstract

Human topoisomerase 1 (TOP1) resolves DNA supercoiling during replication and transcription and is a major target for anticancer therapy. TOP1 poisons exert cytotoxicity by stabilizing the TOP1-DNA cleavage complex (TOP1cc), thereby blocking DNA rejoining and generating lethal DNA damage. Several TOP1 poisons have been approved either as conventional therapeutics or as payloads in targeted delivery systems, and many additional candidates are under clinical development. Here, we resent cryo-EM structures of human TOP1cc bound to eight representative and clinically relevant TOP1 poisons: camptothecin (CPT), six CPT derivatives, and an indenoisoquinoline LMP-400. These cryo-EM structures reveal a TOP1cc conformation that differs substantially from canonical crystal structures. These structures also define how specific modifications on the CPT central scaffold and changing to an alternative non-CPT scaffold reshape drug intercalation geometry, molecular interaction networks, and TOP1cc protein architecture. Together with biochemical trapping data, these structural insights establish a foundation for designing next-generation TOP1 poisons with improved pharmacological properties and for their optimization as antibody-drug conjugate (ADC) payloads.

## Introduction

Human TOP1, a type IB topoisomerase, is a critical enzyme dissipating DNA supercoiling during replication and transcription ^1,2^. It catalyzes a transient cleavage of one DNA strand within the double helix, allowing controlled rotation of the cleaved DNA end to relieve DNA supercoils before religation. This short-lived TOP1-DNA cleavage complex (TOP1cc), containing a DNA nick, is vulnerable to aromatic drug molecules that can intercalate at the DNA break site to trap the complex and prevent DNA religation. All structurally characterized TOP1cc inhibitors exploit this trapping mechanism, acting as TOP1 poisons. They effectively convert TOP1 into a cellular poison that generates lethal DNA damage in rapidly dividing cancer cells ^3^.

The first TOP1 poison, camptothecin (CPT), discovered in the 1970s, is a natural alkaloid isolated from the Chinese tree *Camptotheca acuminata*. Despite its remarkable antitumor activity, the water insolubility, lactone ring instability, and unacceptable toxicity of CPT, precluded its therapeutic use^4,5^. These challenges prompted the development of two parallel strategies: 1) first, synthesis of CPT derivatives with improved water-solubility (topotecan and irinotecan) ^4^ and design of chemically stable non-CPT alternatives^6,7^; 2) more recently, dose-limited toxicity has been improved by targeted delivery systems including liposomes, polyethyleneglycol (PEG) derivatives and, most importantly antibody-drug conjugates (ADCs) to enhance bioavailability and reduce off-target toxicity^3,5^. Several CPT derivatives have achieved clinical approval, including irinotecan and topotecan; liposomal irinotecan; and SN-38 and DXd, as payloads in the ADCs sacituzumab govitecan and trastuzumab deruxtecan, respectively^3^.

Despite these advances, solubility, stability and detailed understanding of how the drugs trap the TOP1cc remain persistent challenges for most candidate drugs and rational design of superior next-generation agents requires detailed structural understanding of how existing drugs bind to and stabilize TOP1ccs. Indeed, only a limited subset of TOP1 poisons has been structurally characterized crystallized in complex with human TOP1 twenty years ago, including topotecan, CPT and three non-CPT indenoisoquinoline compounds^8,9^, limiting our mechanistic understanding of why Dxd and exatecan are more potent than CPT and hampering structure-based drug design of novel TOP1 poisons for further ADC development.

To address this structural knowledge gap and allow further rational drug design, we employed cryo electron microscopy (cryo-EM) for a comprehensive drug binding analysis on seven representative and clinically relevant TOP1 poisons. Here we describe the binding of six CPT derivatives, deruxtecan (Dxd), the payload of two FDA-approved ADCs (Trastuzumab Dxd and Datopotamab Dxd), exatecan, the payload of multiple ADCs in clinical trials, SN-38 the active metabolite of irinotecan and the payload of Sacituzumab govitecan, homocamptothecin (hCPT), two methylenedioxy-camptothecins (MDO-CPTs), and an indenoisoquinoline, indotecan (LMP-400) in clinical trials. We provide molecular insights for understanding TOP1cc stabilization and are consistent with our biochemically measured drug potency. Our findings establish a foundation for the rational design of next-generation TOP1-targeting agents with improved therapeutic properties and for their development as ADC payloads against the most lethal and hard-to-treat tumors.

## Results

### Generating cryo-EM structures of TOP1cc in complex with eight prominent anticancer drugs

To assemble the TOP1cc-drug complexes (Table 1) for cryo-EM analysis, we mixed the functional core of human TOP1 (201-765aa) with a 24-nt dsDNA containing a dominant TOP1 cleavage site (Extended Data Fig.1), in the presence of individual drug molecules. DNA cleavage assays confirmed that all selected compounds efficiently stabilized the TOP1cc (Extended Data Fig.1). Under these sample conditions, we successfully determined cryo-EM structures of TOP1cc bound to exatecan, DXd, hCPT, and two MDO-CPT derivatives at resolutions of 3.2-3.3 Å. However, initial attempts with this setup were unsuccessful for the SN-38 and LMP-400 complexes. Our subsequent sample optimization using full-length TOP1 and a 40-nt dsDNA substrate resolved the two structures at 3.38 Å and 3.08 Å, respectively.

**Table 1.** Cryo-EM data collection, refinement, and validation statistics.

| Table 1. Data collection, processing and model refinement parameters. |  |  |  |  |  |  |  |  |
| --- | --- | --- | --- | --- | --- | --- | --- | --- |
|  | TOP1cc-CPT | TOP1cc-hCPT | TOP1cc-exatecan | TOP1cc-DXd | TOP1cc-SN-38 | TOP1cc-MDO-CPT | TOP1cc-7-CM-MDO-CPT | TOP1cc-LMP-400 |
| PDB code | 10ZS | 10XY | 10WE | 10WD | 10XX | 10WF | 10ZR | 10ZQ |
| EMDB code | EMD-75573 | EMD-75518 | EMD-75501 | EMD-75500 | EMD-75517 | EMD-75502 | EMD-75572 | EMD-75571 |
| **Data collection and processing** |  |  |  |  |  |  |  |  |
| Magnification | 100,000 | 100,000 | 100,000 | 105,000 | 100,000 | 105,000 | 100,000 | 105,000 |
| Pixel size (Å) | 0.830 | 0.830 | 0.830 | 0.832 | 0.830 | 0.832 | 0.830 | 0.824 |
| Defocus range (µm) | -0.8 to -2.4 | -0.8 to -2.4 | -0.8 to -2.4 | -0.8 to -2.4 | -0.8 to -2.4 | -0.8 to -2.4 | -0.8 to -2.4 | -0.8 to -2.4 |
| Voltage (kV) | 200 | 200 | 200 | 300 | 200 | 300 | 200 | 300 |
| Total electron dose (e<sup>-</sup>/Å<sup>2</sup>) | 40 | 46 | 45 | 54 | 45 | 54.2 | 45 | 44 |
| Symmetry imposed | C1 | C1 | C1 | C1 | C1 | C1 | C1 | C1 |
| Particles in reconstruction (no.) | 503k | 517k | 412k | 516k | 510k | 501k | 655k | 510k |
| Map resolution (Å, FSC=0.143) | 3.28 | 3.27 | 3.18 | 3.25 | 3.38 | 3.31 | 3.22 | 3.08 |
| **Model refinement and validation** |  |  |  |  |  |  |  |  |
| Initial model used (PDB code) | 1K4T | 1K4T | 1K4T | 1K4T | 1K4T | 1K4T | 1K4T | 1K4T |
| Mask CC | 0.79 | 0.80 | 0.81 | 0.80 | 0.73 | 0.79 | 0.80 | 0.76 |
| Volume CC | 0.80 | 0.80 | 0.82 | 0.80 | 0.73 | 0.80 | 0.81 | 0.76 |
| Peak CC | 0.76 | 0.77 | 0.78 | 0.75 | 0.69 | 0.75 | 0.78 | 0.70 |
| Model resolution (Å, FSC=0.5) | 3.20 | 3.20 | 3.10 | 3.30 | 3.70 | 3.30 | 3.30 | 3.10 |
| **Average B-factor (Å)** |  |  |  |  |  |  |  |  |
| Protein | 83.48 | 81.92 | 65.15 | 71.30 | 83.60 | 77.44 | 69.22 | 66.28 |
| Nucleotide | 136.40 | 111.47 | 109.13 | 109.38 | 135.62 | 138.72 | 113.58 | 135.95 |
| Ligand | 1.59 | 1.55 | 20.30 | 20.27 | 20.33 | 0.50 | 0.50 | 20.28 |
| **R.m.s. deviations** |  |  |  |  |  |  |  |  |
| RMSD Bond lengths (Å) | 0.005 | 0.004 | 0.005 | 0.006 | 0.005 | 0.006 | 0.004 | 0.005 |
| Bond angles (°) | 0.71 | 0.76 | 0.95 | 0.73 | 0.79 | 0.84 | 0.70 | 0.86 |
| **Ramachandran** |  |  |  |  |  |  |  |  |
| Favored (%) | 96.02 | 94.95 | 95.84 | 95.10 | 94.64 | 94.93 | 95.47 | 95.54 |
| Allowed (%) | 3.98 | 5.05 | 4.16 | 4.90 | 5.36 | 5.07 | 4.53 | 4.46 |
| Outliers (%) | 0.00 | 0.00 | 0.00 | 0.00 | 0.00 | 0.00 | 0.00 | 0.00 |
| **Validation** |  |  |  |  |  |  |  |  |
| Clash score | 6.33 | 6.70 | 4.59 | 6.61 | 11.41 | 7.54 | 5.98 | 8.21 |
| Molprobity score | 1.62 | 1.72 | 1.63 | 2.09 | 1.94 | 1.76 | 1.64 | 1.81 |
| Rotamer outliers (%) | 0.60 | 0.40 | 1.40 | 3.20 | 0.79 | 0.00 | 0.20 | 1.18 |
| C-beta outliers (%) | 0.00 | 0.00 | 0.00 | 0.00 | 0.00 | 0.00 | 0.00 | 0.00 |

Because CPT is the founding TOP1 poison and the parent scaffold for the CPT derivatives analyzed here, it provides an ideal reference for drug binding characterization. To enable direct comparison between CPT and its derivatives, we also determined a cryo-EM structure of TOP1cc-CPT using the same 24-nt dsDNA and TOP1core scaffold used for the CPT-derivative complexes.

### Cryo-EM structures reveal a novel TOP1 complex configuration distinct from canonical crystal structures

Our cryo-EM structure of TOP1cc-CPT immediately revealed significant differences relative to the previously reported TOP1 structures co-crystalized with CPT^9^ (Fig. 1a-c) and other TOP1 poisons^10-12^. In those crystal structures, the dsDNA adopts a compact and straight conformation (Fig. 1c). This rigidity is likely an artifact of crystal packing, driven by the head-to-tail stacking of dsDNA molecules that favors crystallization^12^. In contrast, the cryo-EM structures consistently reveal an approximately 30° bend in the downstream dsDNA, aligning it with the axis of the positively charged linker (Fig. 1b). This conformation reflects a native enzymatic state and further underscores the role of the elongated linker motif in positioning DNA. The downstream DNA is sandwiched between the linker and the nosecone, which both contain multiple positively charged residues in close contact with the DNA phosphate backbone (Fig. 1d-f). The **catalytic domain** (**CAT+CTD**, containing all the catalytic residues)^12,13^ in both the cryo-EM and crystal structures adopt conserved folds, forming an essentially identical catalytic site for DNA backbone positioning and cleavage (Fig. 1d). The linker and the CAP region, especially the nosecone motif, show substantial conformational flexibility. These structural elements readily adjust to changes in DNA conformation. Our cryo-EM structure suggests a partitioned DNA-control mechanism: the catalytic domain and part of the CAP domain form a C-ring that clamps on and secure the upstream dsDNA throughout the catalytic cycle, while the highly positively charged linker and nosecone act as a dynamic clip that sandwiches the flexible downstream DNA and tracks its motion during strand rotation.

**Figure 1.**
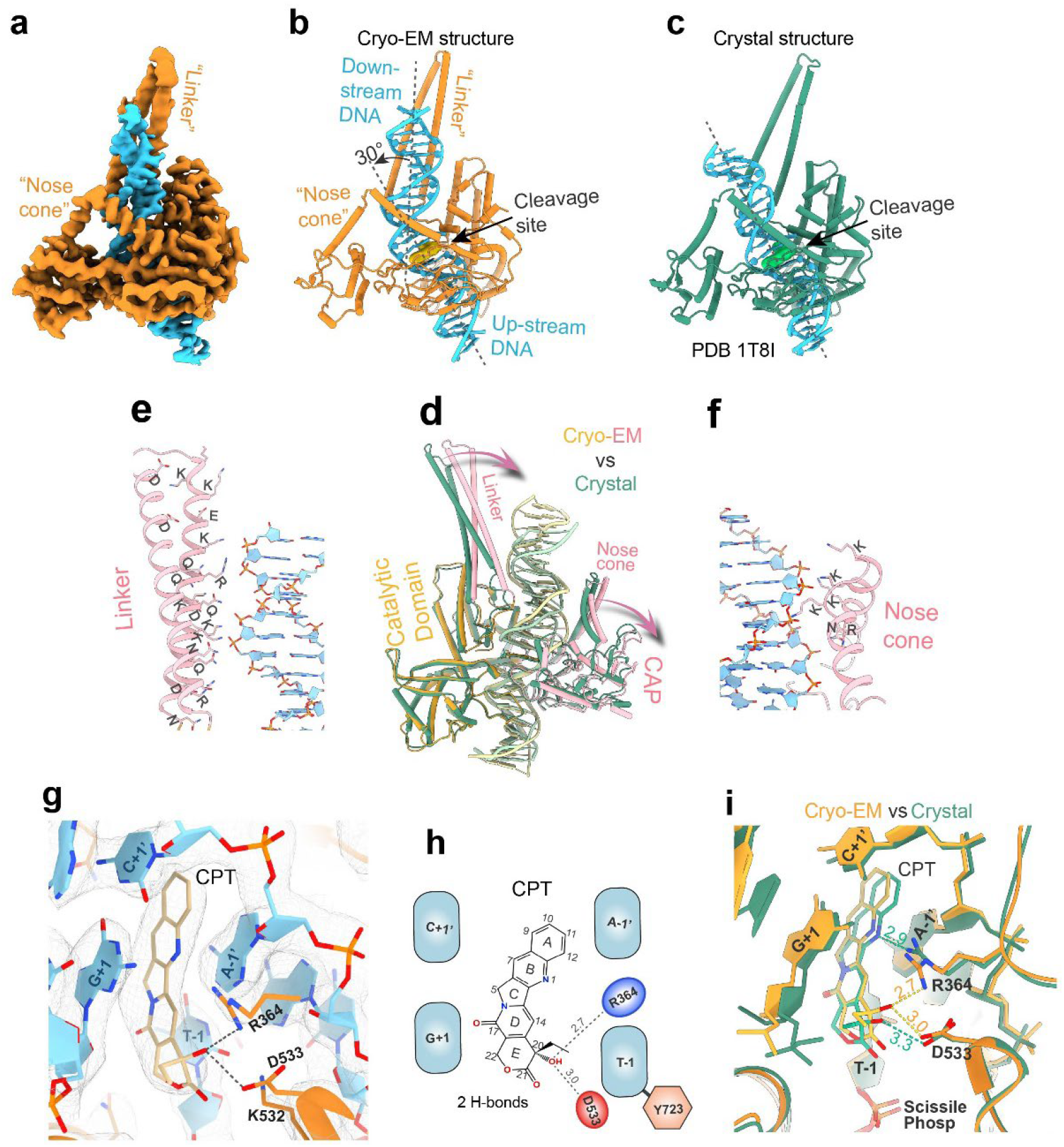
Cryo-EM reveals a distinct TOP1cc configuration compared with canonical crystal structures. **a, b**, Cryo-EM density map and atomic model of TOP1cc in complex with CPT. Key structural elements are highlighted, including the nosecone, linker, and DNA cleavage site with intercalated CPT shown as a yellow surface. **c**, Previously reported crystal structure of the TOP1cc-CPT complex (PDB ID: 1T8I). **d**, Superimposition of the cryo-EM and crystal structures shown in **b** and **c**, aligned on the conserved catalytic domain. The crystal structure is shown in green. In the cryo-EM structure, the catalytic domain is shown in orange, and the linker and CAP domains are shown in light red. Arrows indicate displacement of the linker and CAP domains between the two structures. **e, f**, Detailed views of the linker and nosecone in the cryo-EM structure, highlighting charged residues that either directly interact with DNA or are predicted to contact DNA during DNA rotation. **g**, Cryo-EM density and atomic model of the TOP1cc-CPT active site. Black dashed lines indicate hydrogen bonds. **h**, Chemical structure of CPT, with measured distances corresponding to the interactions shown in **g. i**, Active-site alignment of the cryo-EM and crystal structures, showing CPT displacement and distinct hydrogen-bonding patterns. Dotted lines indicate hydrogen bonds, with distances shown in Å. Interacting protein residues and the four nucleotides at the -1/+1 positions on the cleavage strand and the -1′/+1′ positions on the opposite strand are highlighted.

The cryo-EM map and structure reveal the intercalation of CPT at the DNA nick, stacking against the four DNA bases in an offset parallel configuration (Fig. 1g) and form two hydrogen bonds with R364 and D533 (Fig. 1h). The cryo-EM and crystal structures of TOP1cc-CPT, sharing the four DNA bases at the intercalation site, display similar overall drug-binding geometries (Fig. 1i), while the subtle differences in CPT positioning leads to a distinct drug-protein interaction profile. In both structures, D533 maintains a hydrogen bond with the E-ring hydroxyl group, whereas R364 alternates between interactions with the E-ring hydroxyl and the B-ring amide group (Fig. 1i).

### TOP1 binding by FDA-approved ADC payload SN-38

SN-38 is the active metabolite of the FDA-approved prodrug irinotecan and the payload of the recently FDA-approved ADC sacituzumab govitecan (Trodelvy)^14,15^. In our DNA cleavage assay, SN-38 stabilized TOP1cc more effectively than CPT (Extended Data Fig. 1). To elucidate the molecular basis of this enhanced activity, we compared the cryo-EM structures of TOP1cc bound to SN-38 and CPT.

First of all, SN-38 forms five hydrogen bonds and two van der Waals contacts within the TOP1cc binding site (Fig. 2a,b), establishing a more extensive interaction network than CPT (Fig. 1h). SN-38 binding also produces a slightly narrower DNA gap at the cleavage site (Fig. 2c), suggesting a more compact active-site configuration. This reduced distance between SN-38 and the +1 nucleotide enables two additional interactions with the O5′ and O4′ atoms of the +1 nucleotide (Fig. 2a,b). Moreover, compared with CPT, SN-38 induces tighter DNA clamping by TOP1cc (Fig. 2d). When the conserved TOP1 catalytic domains are aligned, the CAP domains in the two complexes show an RMSD of 1.49 Å (Extended Data Table 1). In addition, a loop within the CAP domain is rearranged in the SN-38-bound structure, positioning R364 closer to the drug (Fig. 2d,e). Consequently, the two terminal amine groups of R364 form two hydrogen bonds with SN-38, one with the B-ring amine and the other with the E-ring hydroxyl group. In contrast, CPT forms only a single hydrogen bond with R364.

**Figure 2.**
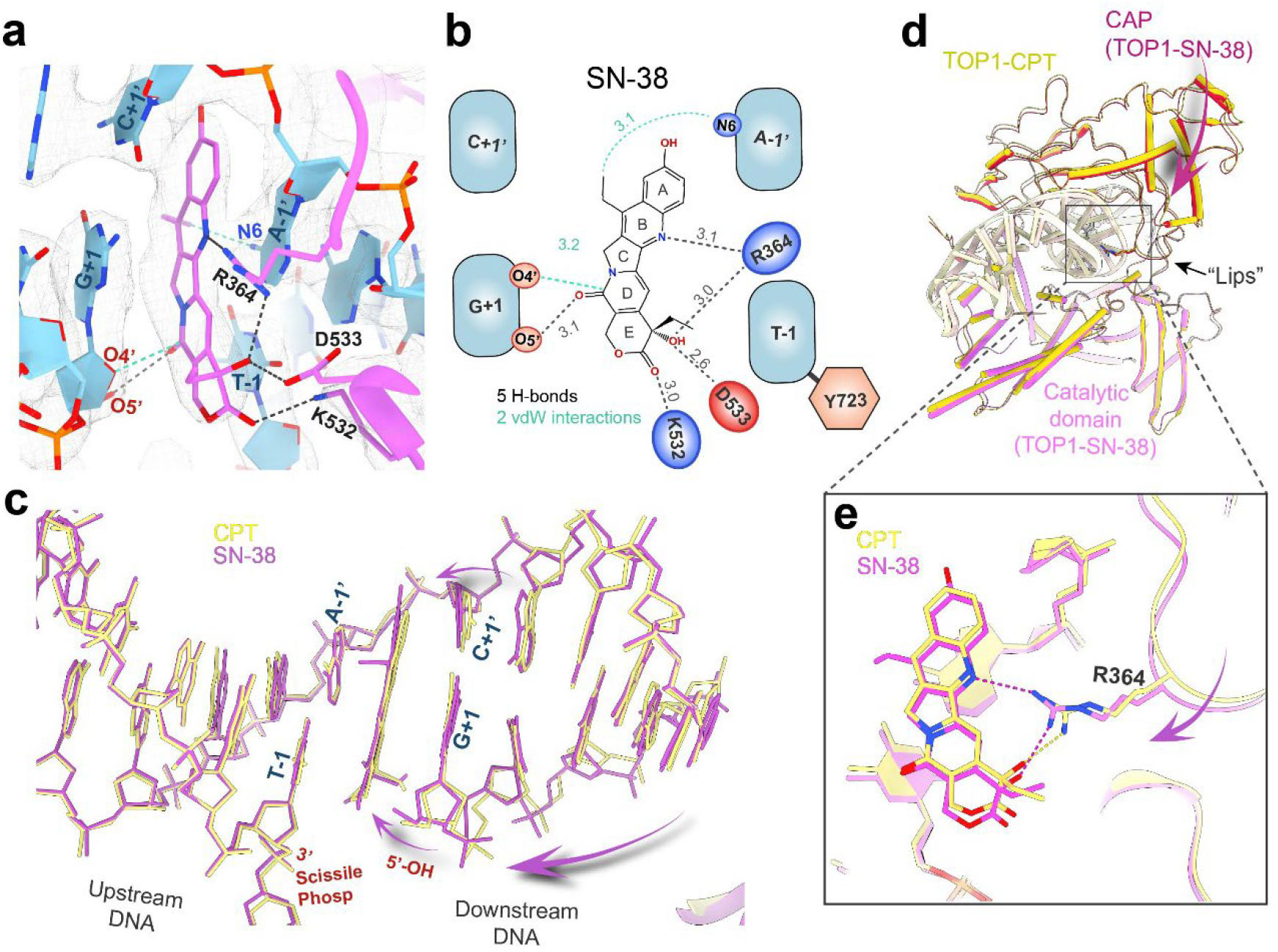
SN-38 binding and comparison with CPT binding. **a**, Cryo-EM density and atomic model of the SN-38 binding site. Interacting protein residues and the four flanking nucleotides are highlighted. Dotted lines indicate hydrogen bonds in black and van der Waals interactions in light cyan. **b**, Chemical structure of SN-38, with corresponding measured distances in **a. c**, Displacement of the downstream DNA bases and phosphate backbone in the SN-38 complex relative to the CPT complex, indicated by arrows. The two structures were superimposed by aligning the TOP1 catalytic domain and enzyme-bound upstream DNA. **d**, Superimposition shows the displacement of the CAP domain in the SN-38 complex in red compared to the CPT complex, demonstrating tighter DNA clamping upon SN-38 binding. The CAP-domain RMSD is 1.49 Å. The whole CPT complex is in yellow; SN-38 complex’s CAP domain is in red and catalytic domain in purple. **e**, Active-site alignment showing repositioning of the R364-containing loop upon SN-38 binding relative to CPT binding, resulting in distinct drug-R364 interactions.

The enhanced binding of SN-38 relative to CPT can be attributed to its chemical modifications on the CPT scaffold: 1) The additional hydroxyl group on the A ring introduces dipolar interactions with the adjacent C+1′ base (Fig. 2a); 2) the B-ring ethyl substituent is electron-rich and likely increases the electron density of the B ring, thereby strengthening its stacking interactions with G+1 and A-1′; 3) the ethyl group forms a dipolar attraction with the 6-amide group of A-1′. Altogether, SN-38’s additional chemical groups lead to enhanced drug interaction network and a more compact TOP1 complex configuration that explain its superior TOP1cc-trapping potency compared to CPT.

### TOP1 binding by ADC payloads exatecan and DXd

Exatecan (DX-8951f) is among the second-generation CPT derivative that exhibit enhanced water solubility, chemical stability and efficacy^5,16^. Although unsuitable as a standalone drug due to unmanageable toxicity, exatecan was successfully developed as an ADC payload and is currently being evaluated in multiple clinical trials^3^. A structurally optimized exatecan variant, DXd, has been successfully integrated into the FDA-approved HER2-targeting ADC trastuzumab deruxtecan (T-Dxd, DS-8201a)^17^. In terms of chemical structure, DXd has replaced a primary amine on exatecan with hydroxyacetamide (Fig. 3a-d) for the ADC assembly.

**Figure 3.**
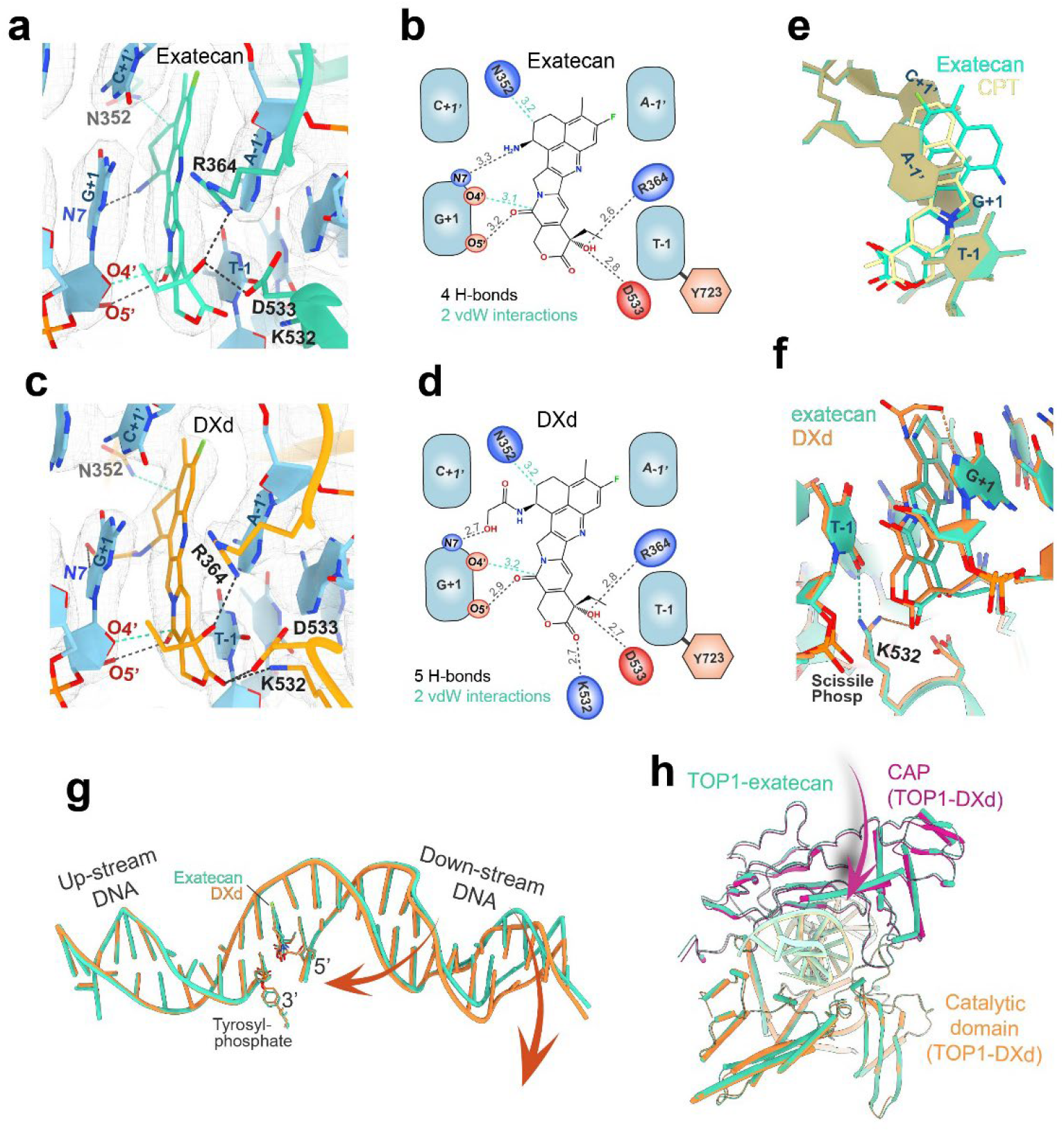
Structural analysis of exatecan- and DXd-bound TOP1cc. **a**, Active-site cryo-EM density and atomic model of the TOP1cc-exatecan complex. Interacting protein residues and the four nucleotides at the -1/+1 positions on the cleavage strand and the -1′/+1′ positions on the opposite strand are highlighted. Dotted lines indicate hydrogen bonds in black and van der Waals interactions in light cyan. **b**, Chemical structure of exatecan, with corresponding distances measured in **a. c, d**, Cryo-EM density, atomic model, and chemical structure of DXd. **e**, Superimposition of the exatecan and CPT intercalation sites, with the four flanking nucleobases labeled. **f**, Superimposition of the exatecan and DXd binding sites. Dotted lines indicate hydrogen bonds formed by exatecan in green and DXd in orange. **g**, Alignment of the TOP1 catalytic domain and upstream DNA from the exatecan and DXd complexes, showing displacement of their downstream DNA. **h**, CAP-domain adjustment upon DXd binding, shown in purple, compared with exatecan binding, shown in green, resulting in a more compact TOP1cc configuration. The CAP-domain RMSD is 1.07 Å. The catalytic domain of the DXd complex is shown in orange and the CAP domain in Magenta; the entire exatecan complex is shown in green.

Both exatecan and DXd outperform CPT in stabilizing TOP1cc (Extended Data Fig. 1). Structural alignment of the active sites in the exatecan- and CPT-bound complexes revealed distinct DNA intercalation poses (Fig. 3e). The altered exatecan binding pose relative to CPT is driven by its addition of a fluorine substituent, a methyl group, and a bulky ring structure, which collectively modify the electron configuration of the CPT scaffold and introduce steric effects. Compared with CPT, exatecan forms two additional hydrogen bonds and two additional van der Waals interactions with the TOP1 complex (Fig. 1g,h vs. Fig. 3a,b). This strengthened binding network accounts for exatecan’s superior TOP1cc stabilization.

DXd is more efficient than exatecan at trapping TOP1cc and appears to be the most potent compound among all tested in our assay (Extended Data Fig. 1). DXd adopts a binding pose similar to exatecan (Fig. 3c,d). The bulky hydroxyacetamide group does not cause significant steric clashes in the binding pocket as it extends toward the empty cavity on the DNA major groove site. We summarize the key differences between DXd and exatecan binding: 1) The hydroxyacetamide terminus of DXd forms a hydrogen bond with the N7 amine of the G+1 base, which is stronger than the corresponding hydrogen bond formed between the amide group of exatecan and the G+1 base (Fig. 3a-d); 2) compared to exatecan, The DXd core scaffold shifts slightly toward the -1 nucleotide position and K532, creating an additional hydrogen bond between its E-ring carbonyl and K532, whereas the K532 interacts with the DNA T-1 base in the exatecan complex instead of the drug (Fig. 3f); 3) DXd binding results in a slightly narrower DNA gap at cleavage site, similar to the SN-38 vs CPT binding, and a greater bending of the downstream DNA backbone (Fig. 3g), which is expected to enhance the alignment between DNA and the linker motif (Fig. 1b); 4) DXd induces a more compact TOP1cc than exatecan, manifested by tighter DNA clamping by the CAP domain (Fig. 3h). When the conserved TOP1 catalytic domains are superimposed, the CAP domains of the DXd and exatecan complexes differ by 1.07 Å RMSD. Relative to the CPT complex, both exatecan and DXd show tighter CAP-domain clamping, with a larger displacement for DXd (RMSD = 1.44 Å) than for exatecan (RMSD = 1.19 Å) (Extended Data Table 1).

Together, the strengthened molecular interactions formed by exatecan and DXd, together with their ability to promote a more compact TOP1cc conformation, account for their enhanced TOP1cc-trapping activity.

### TOP1 binding by representative 10,11-Methylenedioxy-CPT derivatives as ADC payloads

10,11-Methylenedioxy-CPT derivatives (MDO-CPTs) were developed in the 1980s and exhibit substantially higher cytotoxicity than CPT^18,19^. These compounds have also been advanced as ADC payloads and are currently being evaluated in clinical trials^3^ . We analyzed two representative compounds, MDO-CPT and 7-chloromethyl-MDO-CPT, the latter bearing a chloromethyl substitution at the 7-position. Both compounds stabilize TOP1cc more efficiently than CPT (Extended Data Fig. 1).

Our cryo-EM structures show that MDO-CPT and 7-chloromethyl-MDO-CPT adopt nearly identical intercalation poses within the TOP1cc binding site (Fig. 4a,b,e). The two compounds also share a highly similar interaction profile, forming four hydrogen bonds and one van der Waals contact with TOP1cc (Fig. 4a-d). Notably, the electron-rich methylenedioxy ring closely overlaps the C+1′ DNA base at a distance of ∼3 Å (Fig. 4f), anticipated to create strong stacking interactions between them which enhance drug binding affinity. Compared with MDO-CPT, the 7-chloromethyl group of 7-chloromethyl-MDO-CPT appears to have little effect on drug binding, as it extends toward an empty cavity on the DNA major-groove side and does not form strong interactions with TOP1cc (Fig. 4a,b).

**Figure 4.**
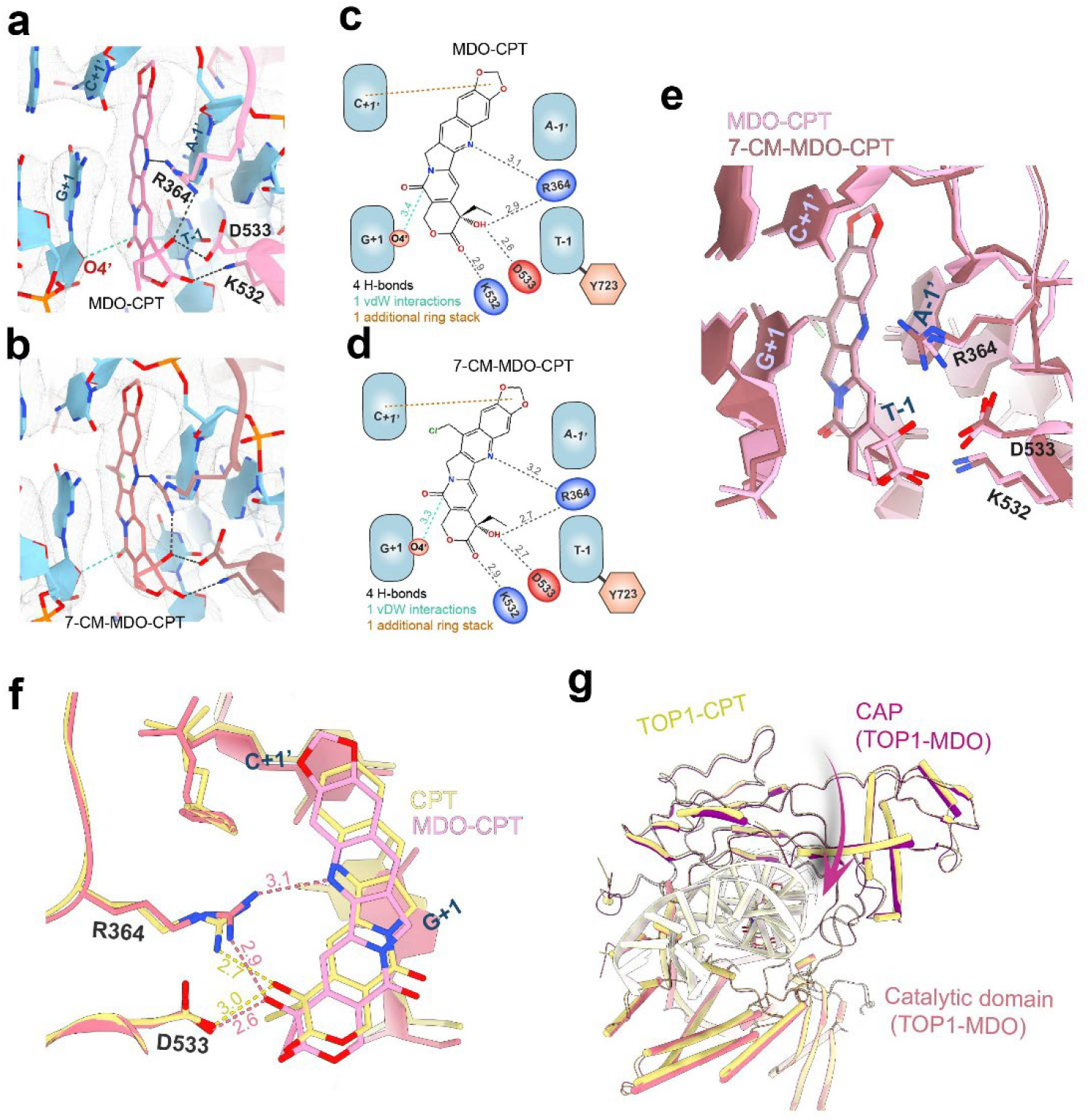
Binding of MDO-CPT and 7-chloromethyl-MDO-CPT and comparison with CPT binding. **a, b**, Cryo-EM density and atomic models of TOP1cc in complex with MDO-CPT and 7-chloromethyl-MDO-CPT. Interacting protein residues and the four active-site nucleotides are highlighted. Dashed lines indicate hydrogen bonds in black and van der Waals contacts in light cyan. **c, d**, Chemical structures of MDO-CPT and 7-chloromethyl-MDO-CPT, with measured distances corresponding to the interactions shown in **a** and **b. e**, Superimposition of the drug-binding sites, showing comparable intercalation conformations for the two CPT analogs. The four catalytic-center nucleotides and protein residues involved in ligand interactions are highlighted. **f**, Active-site alignment showing binding poses and interactions of MDO-CPT and CPT. **g**, Structural comparison of the CAP domain in the MDO-CPT-bound complex, shown in dark red, and the CPT-bound complex, shown in yellow, demonstrating enhanced DNA clamping upon MDO-CPT binding. The CAP-domain RMSD is 1.22 Å. The CPT complex and the domains of MDO-CPT complex are colored as indicated.

Like the potent CPT derivatives discussed above, the MDO-CPTs also induce tighter TOP1-mediated DNA clamping than CPT (Fig. 4g and Extended Data Table 1), indicating formation of a more compact and stable TOP1cc-drug complex.

### TOP1 binding by Homocamptothecin, an E-ring modified CPT and promising clinical candidate

In physiological solution, the E-ring of CPT exists in equilibrium between closed (active) and open (inactive) configurations^20^. Homocamptothecin (hCPT), an E-ring-modified CPT derivative, was developed to improve chemical stability^21^. Beyond its enhanced chemical stability, hCPT also exhibits superior TOP1cc trapping activity (Extended Data Fig.1).

When bound to TOP1cc, the modified seven-membered E-ring of hCPT is oriented nearly perpendicular to the drug central plane, whereas the classical lactone ring of CPT lies approximately parallel (Fig. 5a,c). When hCPT intercalates into the cleavage site, the perpendicular orientation of the E-ring carbonyl group creates steric repulsion with the 2-carbonyl of base T-1, likely driving the observed shift in drug position relative to CPT (Fig. 5c). Consequentially, this repositioning generates additional drug interactions (Fig. 5a,b vs Fig. 1h): 1) a new hydrogen bond between K532 and the E-ring carbonyl; 2) a hydrogen bond between R364 and the B-ring amine; 3) a hydrogen bond and a van der Waals contact between the D-ring carbonyl and the 5′-OH and the 4′-ether of the +1 nucleotide. This expanded interaction network enhances binding enthalpy and affinity, providing a structural explanation for the increased potency of hCPT in trapping TOP1cc observed in our biochemical assay. Despite the differences in active-site interactions, the overall domain architectures of the CPT- and hCPT-bound complexes are nearly identical (Fig. 5d).

**Figure 5.**
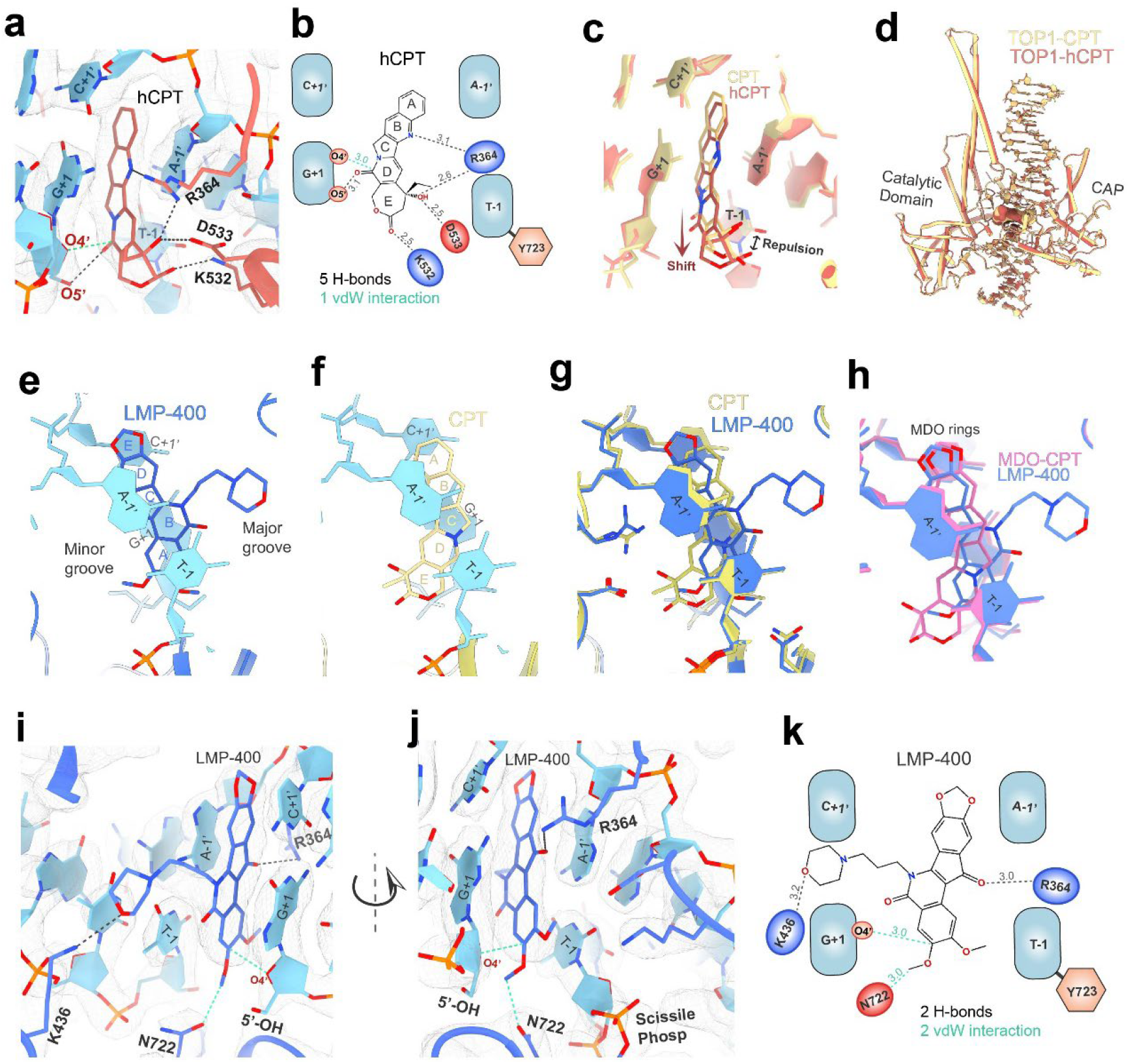
TOP1cc binding by hCPT and LMP-400, compared with CPT binding. **a, b**, Cryo-EM density and structural models of the hCPT binding site and molecular interactions. Black dotted lines indicate hydrogen bonds, and light cyan lines indicate van der Waals interactions. Measured distances are shown in Å. **c**, Superimposition of the CPT and hCPT intercalation sites, with the four flanking nucleobases shown. **d**, Superimposition of CPT- and hCPT-bound TOP1cc structures, showing similar overall protein configurations. **e, f**, LMP-400 and CPT binding sites shown from the same viewing angle, comparing their drug intercalation poses and DNA base-stacking profiles. The five rings of each compound are numbered according to the literature. **g, h**, Superimposition of drug-binding sites comparing LMP-400 with CPT and LMP-400 with MDO-CPT. **i, j**, Cryo-EM density maps and corresponding atomic models of the LMP-400 binding site, shown from two viewing angles. Key interacting protein residues and the four active-site nucleotides are highlighted. Black dashed lines indicate hydrogen bonds, and light cyan dashed lines indicate van der Waals interactions. **k**, Chemical structure of LMP-400, with corresponding distances measured in **i** and **j**.

These structural and biochemical features support hCPT as a promising candidate scaffold for ADC payload development.

### TOP1 binding by LMP-400, a representative non-CPT compound

To date, all FDA-approved anticancer drugs that target TOP1, either as standalone agents or as ADC payloads, are CPT derivatives. Most TOP1-targeting drugs currently in clinical development are also based on the CPT scaffold. Non-CPT TOP1 poisons have also been developed. A representative example is LMP-400 (indotecan), one of the indenoisoquinolines^22^ that has currently been evaluated in clinical trials by the National Cancer Institute.

In our cryo-EM structure, the unique five-ring system of LMP-400 distinguishes it from CPT derivatives and produces a distinct DNA-stacking profile (Fig. 5e,g). The molecule is nearly planar, except for a slightly bent methylenedioxy ring. All five rings of LMP-400 exhibit displaced-parallel π-π stacking with surrounding DNA bases, with increased DNA base overlap compared to CPT (Fig. 5e-g). In contrast, only the A, B, and C rings of CPT significantly contribute to DNA base stacking. Interestingly, the MDO rings of both LMP-400 and MDO-CPT occupy similar positions relative to the DNA C+1’ base (Fig. 5h).

LMP-400, along with LMP-776 and LMP-744 (another two indenoisoquinolines in clinical trials at NCI), features a flexible long arm attached to the B-ring, with LMP-400 showing the strongest TOP1cc trapping (Extended Data Fig. 2). In our cryo-EM structure, the LMP-400 arm extends into a large cavity facing the DNA major groove (Fig. 5e), improving shape complementarity between the drug and the binding pocket. The terminal ether of LMP-400 lies within hydrogen-bonding distance of K436 (Fig. 5i). In addition, the central scaffold of LMP-400 forms one hydrogen bond and two van der Waals interactions with TOP1cc (Fig. 5i-k).

Overall, LMP-400 exhibits enhanced DNA stacking and improved shape complementarity with the binding pocket but forms weaker and/or fewer hydrogen bonds than CPT and derivatives (Extended Data Table 1). Therefore, its binding is driven more by entropy than enthalpy. In our biochemical tests, LMP-400 is less effective at stabilizing TOP1cc than those CPT derivatives which consistently establish stronger hydrogen bond networks (Extended Data Fig. 1). These results highlight the importance of strong molecular interactions for achieving maximal drug-mediated TOP1cc stabilization.

## Discussion

Our cryo-EM study reveals how CPT, six CPT derivatives, and a representative non-CPT analog bind human TOP1cc. All eight compounds intercalate between the four nucleotides flanking the TOP1-mediated DNA cleavage site. Despite sharing this general trapping mechanism, each compound establishes a specific molecular interaction profile, adopts a distinct intercalation geometry, and induces drug-dependent structural remodeling of TOP1cc.

Comparative structural analysis reveals how differential drug binding and induced conformational rearrangements correlate with the drug efficacies observed in our biochemical assays. First, a greater number of hydrogen bonds contributes to stronger TOP1cc trapping, as illustrated by the six CPT derivatives, which showed greater efficacy than CPT and LMP-400 (Extended Data Table 1a). Second, enhanced drug binding and potency are often associated with tighter DNA clamping, as observed in the DXd, SN-38, exatecan, and MDO-CPT complexes compared with the CPT complex (Extended Data Table 1b). Although LMP-400 also induces tighter DNA clamping, its relatively weak TOP1cc-trapping activity is likely explained by its less extensive hydrogen-bonding network. Together, these findings suggest that potent next-generation TOP1cc-targeting compounds should consider chemical modifications that can strengthen a hydrogen-bonding network while preserving favorable DNA stacking and pocket complementarity.

Our cryo-EM structures also reveal substantial unoccupied cavities near the bound drugs (Extended Data Fig. 3). These spaces could be exploited by introducing appropriate chemical groups onto drug scaffolds to improve pocket complementarity and enhance binding affinity. They also provide suitable positions for introducing chemical handles required for ADC conjugation while minimizing steric clashes. Several compounds analyzed here already benefit from such scaffold extensions. For example, the extended arms of LMP-400 and DXd partially occupy a cavity on the DNA major-groove side, whereas the additional methylenedioxy rings of MDO-CPTs and LMP-400 partially fill a distal cavity and improve stacking with DNA bases.

## Acknowledgements

Cryo-EM data were collected at the Center for Structural Biology cryo-EM facility (NCI Frederick), NICE-NIH Intramural Cryo-EM Consortium, and the MICEF Cryo-EM Facility. We thank Dan Shi, Mi Li, Rick Huang, Haotian Lei, Yanxiang Cui, Huaibin Wang, and Ulrich Baxa for assistance with cryo-EM data collection. This work was supported by the Center for Cancer Research, NIH (Z01-BC006161) to Y.P. and by the Fondation pour la Recherche Médicale (AJE202512051057) to X.Y.

---

**Extended Data Table 1.** Summary of drug-TOP1cc interactions and CAP-domain displacement. **a**, Summary of drug-TOP1cc interactions, including the number of hydrogen bonds, van der Waals contacts, and other observed significant interactions, excluding the stacking interactions between DNA bases and the CPT/indenoisoquinoline core scaffold. **b**, Pairwise comparison of CAP-domain displacement. RMSD values between CAP domains were calculated after aligning the conserved TOP1 catalytic domains and enzyme-bound upstream DNA. A higher RMSD indicates that the latter drug complex induces tighter DNA clamping than the former.

| Drug complex | # of H-bond | # of vdW | Additional strong interactions |
| --- | --- | --- | --- |
| CPT | 2 | 0 |  |
| hCPT | 5 | 1 |  |
| exatecan | 4 | 2 |  |
| DXd | 5 | 2 |  |
| SN-38 | 5 | 2 |  |
| MDO-CPT | 4 | 1 | MDO ring & C+1' base |
| 7-CM-MDO-CPT | 4 | 1 | MDO ring & C+1' base |
| LMP-400 | 2 | 2 |  |

**Extended Data Table 1. Summary of drug-TOP1cc interactions and CAP-domain displacement**
| RMSD of the CAP domains between the indicated drug-TOP1cc complexes |  |
| --- | --- |
| CPT vs. hCPT | 0.95 |
| CPT vs. exatecan | 1.19 |
| CPT vs. DXd | 1.44 |
| CPT vs. SN-38 | 1.49 |
| CPT vs. MDO-CPT | 1.22 |
| CPT vs. 7-CM-MDO-CPT | 1.1 |
| CPT vs. LMP-400 | 1.45 |
| Exatecan vs. DXd | 1.07 |

**Extended Data Figure 1.**
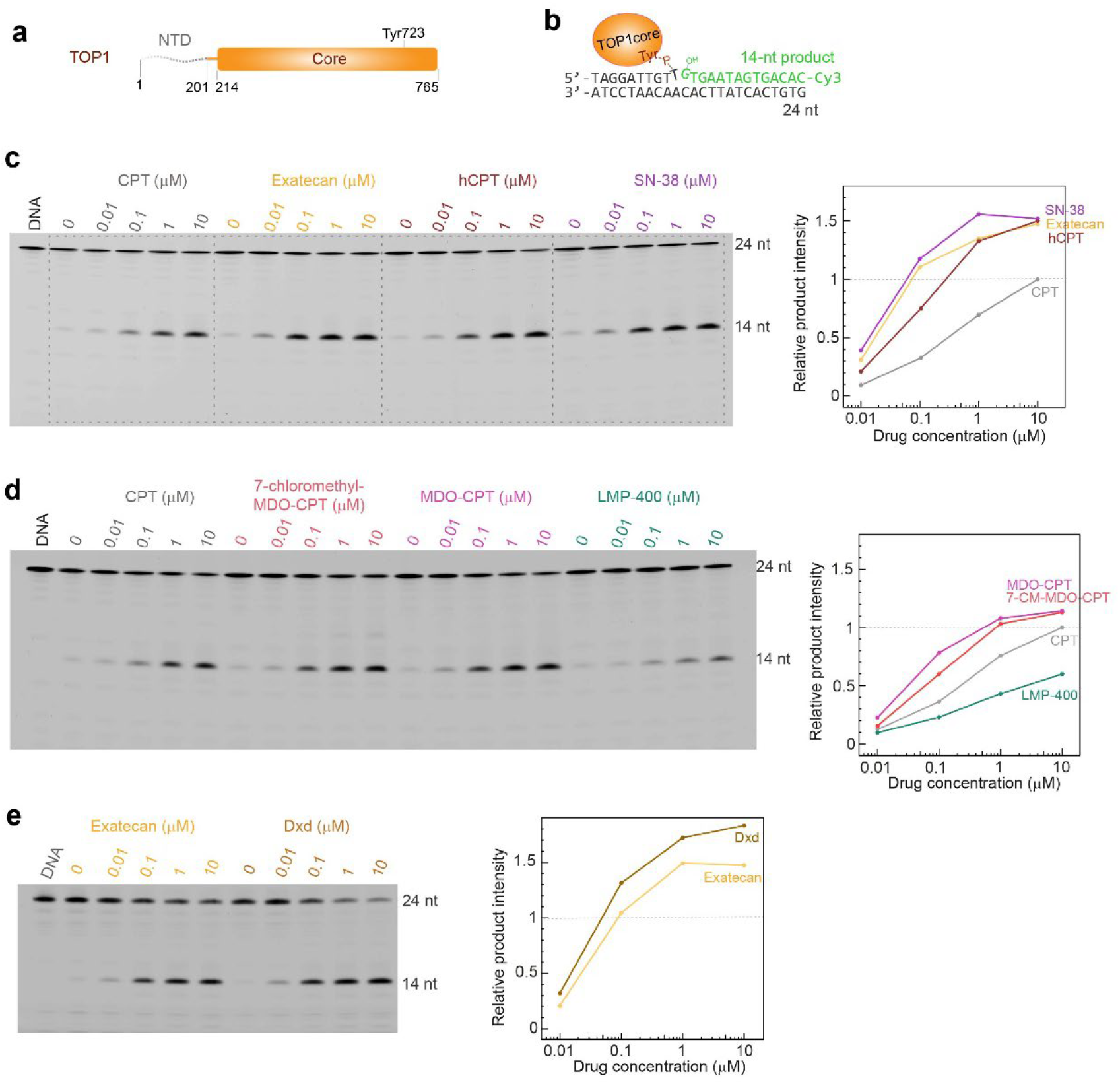
Human TOP1 domain organization, DNA substrate design, and drug-dependent TOP1cc stabilization. **a**, Schematic of human TOP1 domain architecture, including the structurally flexible N-terminal domain (NTD; residues 1-201) and the conserved core domain (residues 214-765). The catalytic tyrosine residue Tyr723 is highlighted. **b**, The 24-bp DNA substrate used for both DNA cleavage assays and cryo-EM analysis. TOP1-mediated cleavage generates a covalent bond between Tyr723 and the scissile phosphate of DNA. The Cy3-labeled cleavage strand, corresponding to the 24-nt substrate, and the resulting Cy3-labeled 14-nt cleavage product can be resolved and detected by denaturing urea-PAGE. **c-e**, TOP1 DNA cleavage assays performed with the indicated drugs and concentrations, with corresponding urea-PAGE gels shown. The 14-nt cleavage products were quantified and plotted in the graphs on the right. The y axis represents the relative amount of 14-nt DNA product, normalized to the band intensity observed with 10 μM CPT.

**Extended Data Figure 2.**
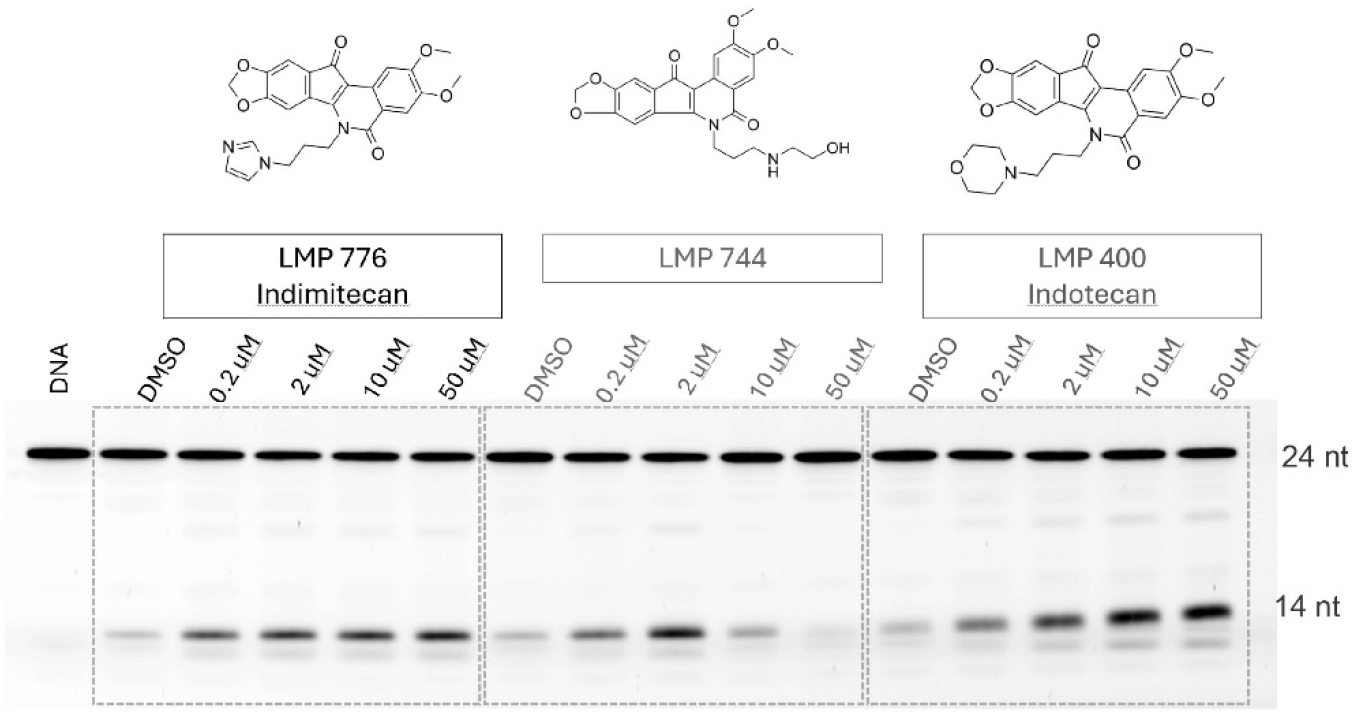
TOP1 DNA cleavage assays with indenoisoquinoline derivatives. TOP1 DNA cleavage assays performed with three indenoisoquinoline derivatives at four concentrations. The 24-nt band corresponds to the intact DNA substrate, whereas the 14-nt band represents the cleavage product.

**Extended Data Figure 3.**
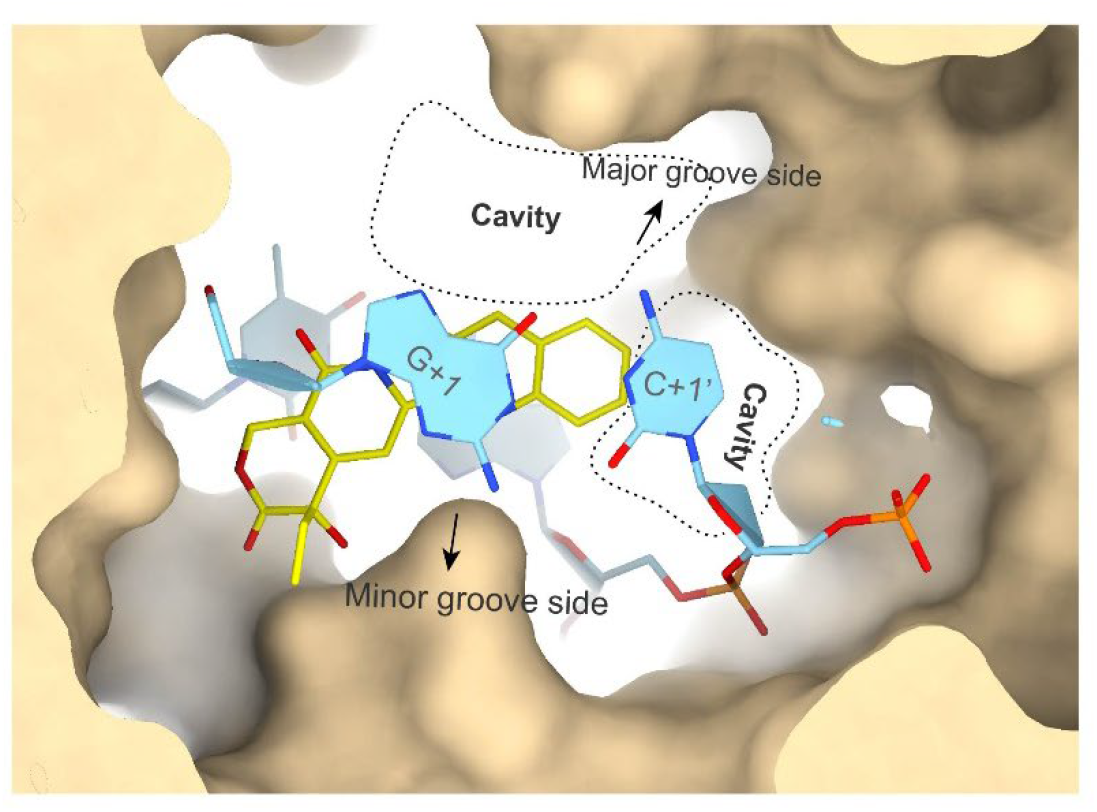
Empty cavities in the CPT-bound TOP1cc active site. Active site of CPT-bound TOP1cc, showing empty cavities on the DNA major-groove side and beyond the CPT A ring, indicated by dotted circles.

**Supplementary Information Figure 1.**
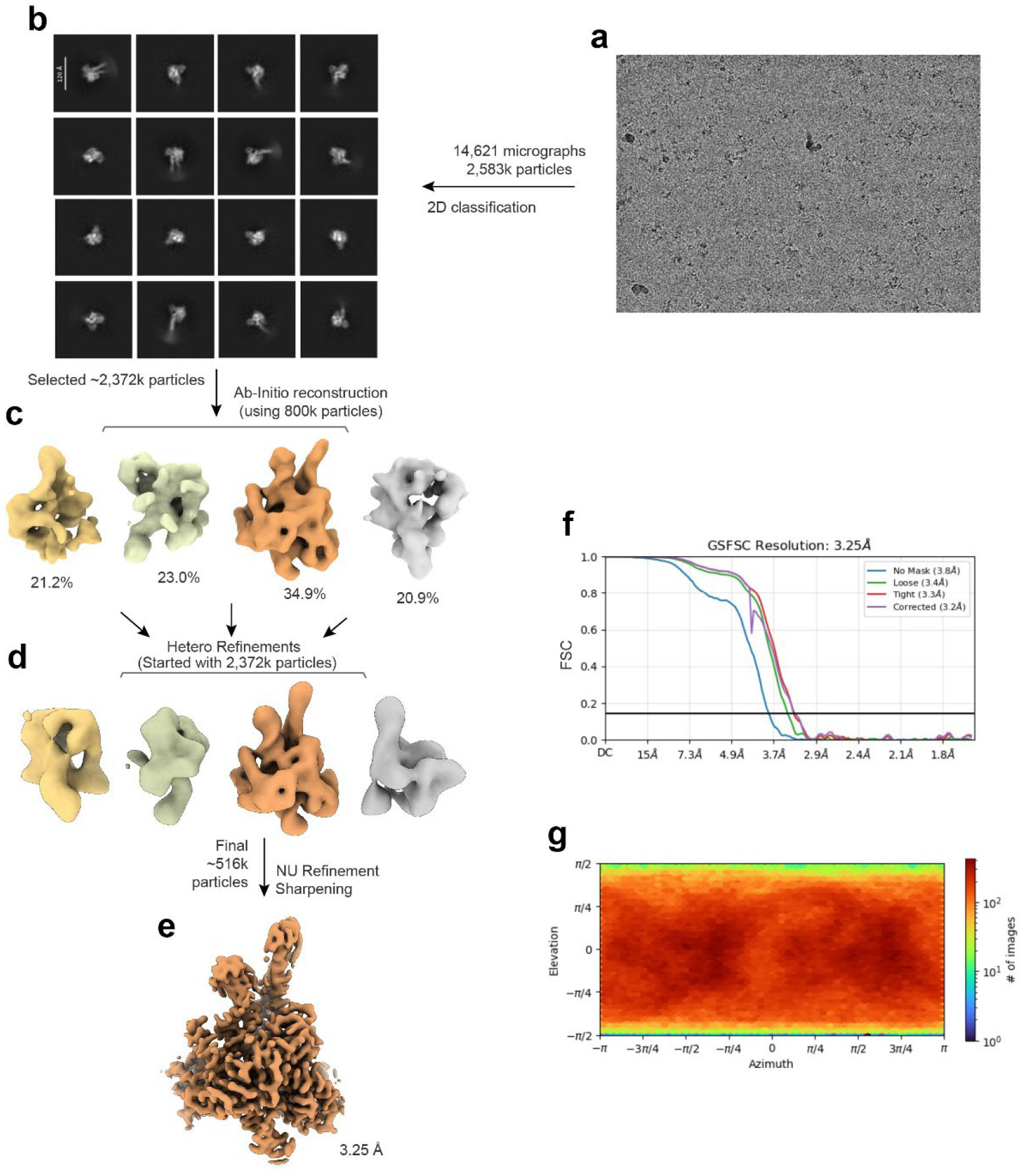
Cryo-EM structure determination of the DXd-bound TOP1 DNA cleavage complex. **a-e**, Cryo-EM data-processing workflow for the TOP1cc-DXd complex in cryoSPARC. This workflow is representative of the procedures used for all cryo-EM maps presented in the manuscript. **f**, Fourier shell correlation (FSC) curve for the final map, calculated after the map refinement shown in **e**. The gold-standard FSC 0.143 cutoff is indicated. **g**, Heatmap showing the angular distribution of particles contributing to the final reconstruction.

